# Automated control of odor dynamics for neurophysiology and behavior

**DOI:** 10.1101/2021.08.17.456668

**Authors:** Luis Hernandez-Nunez, Aravinthan D. T. Samuel

**Affiliations:** Department of Physics, Harvard University, Cambridge, MA 02138, USA; Center for Brain Science, Harvard University, Cambridge, MA 02138, USA; Systems, Synthetic, and Quantitative Biology PhD program, Harvard University, Cambridge, United States

## Abstract

Animals use their olfactory systems to avoid predators, forage for food, and identify mates. Olfactory systems detect and distinguish odors by responding to the concentration, temporal dynamics, and identities of odorant molecules. Studying the temporal neural processing of odors carried in air has been difficult because of the inherent challenge in precisely controlling odorized airflows over time. Odorized airflows interact with surfaces and other air currents, leading to a complex transformation from the odorized airflow that is desired to the olfactory stimulus that is delivered. Here, we present a method that achieves precise and automated control of the amplitude, baseline, and temporal structure of olfactory stimuli. We use this technique to analyze the temporal processing of olfactory stimuli in the early olfactory circuits and navigational behavior of larval *Drosophila*. Precise odor control and calcium measurements in the axon terminal of an Olfactory Receptor Neuron (ORN-Or42b) revealed dynamic adaptation properties: as in vertebrate photoreceptor neurons, Or42b-ORNs display simultaneous gain-suppression and speedup of their neural response. Furthermore, we found that ORN sensitivity to changes in odor concentration decreases with odor background, but the sensitivity to odor contrast is invariant – this causes odor-evoked ORN activity to follow the Weber-Fechner Law. Using precise olfactory stimulus control with freely-moving animals, we uncovered correlations between the temporal dynamics of larval navigation motor programs and the neural response dynamics of second-order olfactory neurons. The correspondence between neural and behavioral dynamics highlights the potential of precise odor temporal dynamics control in dissecting the sensorimotor circuits for olfactory behaviors.

## Introduction

Olfactory systems overcome two types of detection challenges. One, because natural odors are chemically diverse and typically occur in mixtures, olfactory systems solve a combinatorial problem to determine chemical identity. Two, because odors are often carried in dynamic airstreams, olfactory systems also decode the temporal structure of olfactory stimuli. While the problem of the combinatorial olfactory code is widely studied (***Pashkovski et al., 2020***; ***Parnas et al., 2013***; ***Mathew et al., 2013***; ***Si et al., 2019***), we know less about how the temporal dynamics of olfactory stimuli are encoded in neural circuits and behavior.

Precise control of the temporal dynamics of visual and auditory stimuli has revealed computationally-relevant response properties in the neural dynamics of photoreceptors and hair cells (***Hosoya et al., 2005***; ***Laughlin, 1989***; ***Frank et al., 1999***). Conducting similarly precise experiments is difficult when studying the olfactory system because odorized airflows are hard to control over time. Odorized airflows can have complex and unpredictable interactions with surfaces and other airflows, complicating the relationship between the flow of odorized air that is programmed and the olfactory stimulus that is delivered. For this reason, most studies of olfactory systems use simple odorized air puffs, but even air puffs can suffer from trial-to-trial variability (***Gorur-Shandilya et al*.** (***2019***)).

In small animals that swim or burrow through olfactory environments – such as *C. elegans* and larval *Drosophila* – the problem of precise stimulus control of airflow can be circumvented by using microfluidics to deliver odors carried in water (***Si et al., 2019***; ***Kato et al., 2014***; ***Van Giesen et al., 2016***). However, the laminar nature of water flows in microfluidic channels prevents these devices from continuously modulating the concentration of odor over the time course of stimulus delivery. As a result, the stimuli that can be generated is limited to trains of rectangular odor pulses controlled by ON/OFF valves. Additionally, microfluidics do not necessarily recapitulate the olfactory experience of navigating odor landscapes in air.

Depending on the wind and spatial scale, odor landscapes in air can be dominated by diffusion, convection, or turbulence. In *Drosophila*, navigational behavior and Olfactory Receptor Neuron (ORN) responses in turbulent odor landscapes have been shown to depend on the statistics of odor encounters (***Gorur-Shandilya et al., 2017***; ***Demir et al., 2020***). In diffusion-dominated and laminar odor landscapes, odor concentrations vary smoothly; in this regime, behavioral and neural responses have been shown to depend on the temporal derivative of odor concentration (***Schulze et al., 2015***; ***Gershow et al., 2012***). While it is possible to study animal behavior in turbulent odor landscapes (***Demir et al., 2020***), being able to precisely manipulate the temporal properties of smooth odor waveforms to dissect neural and behavioral responses remains challenging. One method is to use feedback loops to control odorized airflows – e.g., opening and closing solenoid valves or massflow controllers with proportional, integral, and derivative (PID) control mechanisms (***Mathew et al., 2013***; ***Gorur-Shandilya et al., 2019***; ***Schulze et al., 2015***). However, while these techniques control the flow of odorized air at specific points within the odor delivery apparatus, they do not deal with the subsequent, unavoidable, and unpredictable airflow perturbations that occur before the olfactory stimulus reaches the animal.

Here, we present a control strategy that compensates for perturbations and enables precise control of the dynamics of odor waveforms. First, we quantify the performance limitations of Proportional, Integral, and Derivative (PID) massflow controllers that deliver odorized airflows. Next, we build a *non-linear autoregressive with exogenous input* (*NARX*) model to capture the dynamics of the overall odor delivery system. Using the NARX model we design a second PID controller that, in tandem with the massflow PID controller, is able to cancel perturbations and achieve precise control of the frequency, amplitude, and baseline of temporal odor stimuli.

We use our method to study neural dynamics in the early olfactory system of larval *Drosophila*. By analyzing ORN calcium responses to olfactory stimuli of identical shape and baseline but different amplitude, we discover a case of non-linear dynamic adaptation. Gain suppression in Or42b-ORN is accompanied by a speedup of its neural response. Using stimuli of identical shape and amplitude but different baseline, we find that Or42b sensitivity diminishes as odor baseline increases. Furthermore, using stimuli of identical contrast and shape but different baseline, we find that the O42b-ORN neural response follows Weber’s Law such that its sensitivity to stimulus contrast is independent of odor baseline.

We extend our method to analyzing the navigational behavior of freely crawling larvae exposed to a laminar flow of odorized air. With our setup, we were able to map the temporal parameters of the odor stimulus to navigational dynamics. By comparing the responses of larval *Drosophila* to calibrated stimuli in our physiology experiments and our behavior experiments, we identify dynamical features of behavior that directly reflect second-order olfactory neuron dynamics.

## Results

### The challenges of odor temporal control

Typical closed-loop odor delivery systems separately modulate clean and odorized airflows. The two air streams then converge to induce temporal changes in the odor concentration of the mixed airflow(***Gorur-Shandilya et al., 2017***; ***Schulze et al., 2015***). Here, we also use two massflow controllers (MFCs) to separately control the amount of clean air and the amount of odorized air that are mixed in a Y-junction before being delivered to the animal. In the odorized air branch, one MFC outlet is connected to a muffler that produces bubbles inside a bottle containing an odor-water dilution. Bubbling produces odor vapor which flows out through another port in the bottle cap. The odorized airflow enters a solenoid shuttle valve – when the solenoid is OFF the odorized airflow is disposed through a vacuum port; when the solenoid is ON the odorized airflow enters the Y-junction where it mixes with the clean airflow. A photoionization detector measures odor concentration at the output of the odor delivery system (Fig. 1A, Fig.1-Supp.Fig. 1A,B).

**Figure 1.**
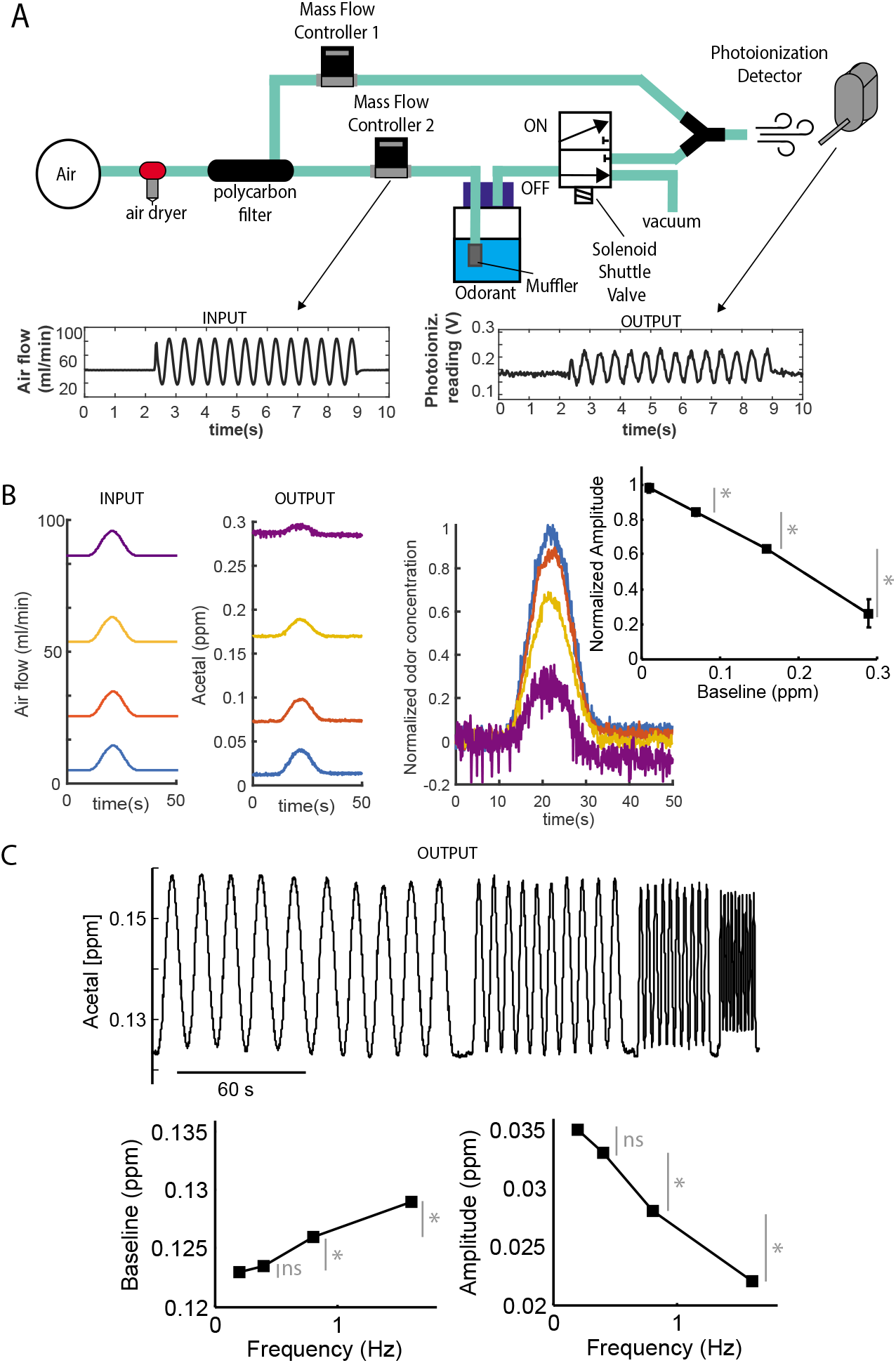
The challenges of odor temporal control. **(A)** Schematic representation of the odor delivery system. Clean air is successively dried and filtered before entering two separate Mass Flow Controllers (MFCs). MFC-1 controls the amount of clean air that is then mixed with odorized air before been delivered. MFC-2 controls the amount of air that enters a small bottle through a muffler that generates bubbles. Bubbling inside the bottle then generates odor vapors that flow out of the bottle through a second port. Odorized air then passes through a shuttle valve and finally mixes with the clean air to generate a temporal odor waveform. The input temporal signal is applied to MFC-2, while MFC-1 simply compensates for the input to MFC-2 to keep the overall airflow constant. The output is the odor concentration measured with a photoionization detector at the output of the odor delivery system. **(B)** Inputs of identical shape and amplitude (peak minus baseline) result in outputs of decreasing amplitude. The normalized odor concentration is each output odor concentration waveform minus its initial concentration at t=0. The output amplitude at each successive baseline is significantly different than the previous one (p<0.005 with Wilcoxon matched pairs test). **(C)** Using sinusoidal waves of different frequencies but identical amplitude and baseline as input, results in an odor concentration waveform with increasing baseline and decreasing amplitude. Baseline is the average of the local minima of each sinusoidal wave. Grey asterisks and lines denote significantly different values of baseline or amplitude between sinusoidal waves of different frequency (p<0.005 with Wilcoxon matched pairs test). **Figure 1–Figure supplement 1.** Figure 1–Figure Supplement 1. Olfactometer components (**A**) and graphical user interface (GUI) (**B**).

The input waveform to the MFC in the odorized air branch results in an output odor concentration waveform that is measured with the photoionization detector. For example, a sine wave of airflow generated by the MFC approximately produces a sine wave of odor concentration (Fig. 1A insets). However, regardless of the parameter selection in the MFCs, the transformation between the input waveform of odorized airflow and the output waveform of odor concentration is not linear. This transformation is perturbed by interactions between the clean and odorized airflows, and interactions with the muffler, the valve, and other surfaces.

The non-linear transformation from the stimulus that is programmed to the stimulus that is delivered is evident, for example, when comparing odor waveforms with the same shape but different baselines (background odor concentration). Inputs that differ only in baseline produce outputs that, in addition, differ in amplitude (Fig. 1B left). As the baseline increases, the amplitude of the odor waveform decreases (Fig. 1B right). Similarly, sinusoidal inputs of identical baseline and amplitude but different frequency produce outputs that also differ in amplitude and baseline (Fig. 1C). As the frequency increases, the amplitude of the sinusoidal stimulus decreases and the baseline increases (Fig. 1C).

### Modelling an odor delivery system

In order to precisely control the baseline, frequency, and amplitude of an odor waveform, we need to actively compensate for multiple non-linear effects within our apparatus. To do this, we derived a mathematical model that captures the odor delivery system input-output performance. We then used this mathematical model to design a controller that enables compensating for the inherent system non-linearities.

The operation of our odor delivery system may be viewed as a series of functional blocks (Fig. 2A). The desired stimulus or set-point is the input to the MFC. The MFC has an internal loop that consists of a Proportional, Integral and Derivative (PID) controller that drives flow modulation actuators, using feedback from a flow sensor to regulate the airflow. The difference between the measured airflow and the desired airflow is the input to the PID controller, which updates the input to the actuators to minimize this difference. The output of the MFC is perturbed by many different interactions within the setup before it reaches the animal – e.g., interactions in the muffler and odor dilution bottle, and additional perturbations when the clean and odorized airflows are mixed. We sought to model the complete transformation from the set-point that is communicated to the MFC to the actual waveform of olfactory stimulus (Fig. 2A).

**Figure 2.**
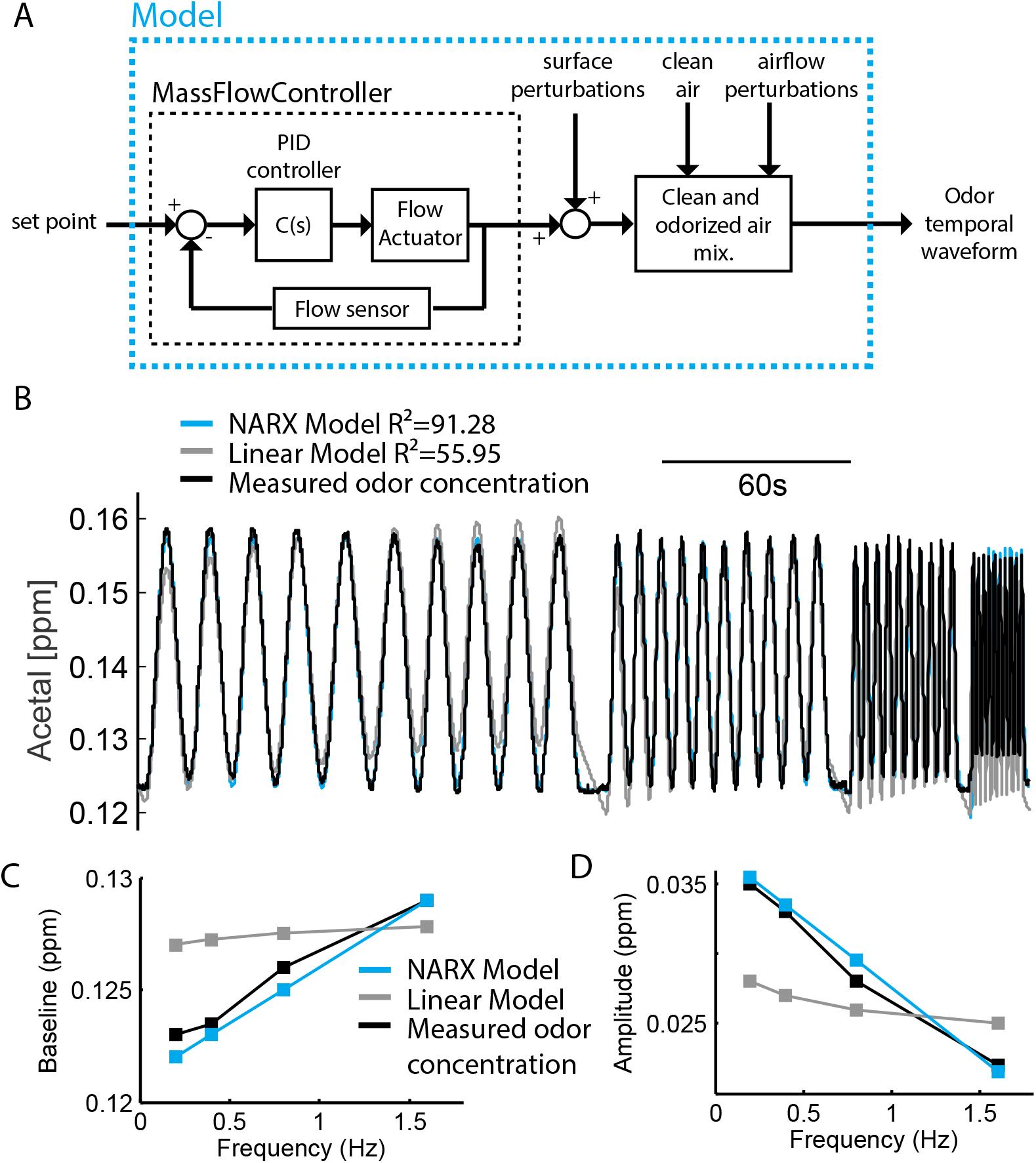
Systems identification of an olfactometer. **(A)** Transfer-function diagram of the olfactometer. The Mass Flow Controller (MFC) includes a ‘Flow Actuator’ that modulates flow and it is controlled by a Proportional, Integral, and Derivative (PID) Controller. The PID controller tries to minimize the error between the programmed set-point and the flow measured by a ‘Flow Sensor’. The output of the MFC then receives surface perturbations as it travels in a tubing and after exiting the muffler inside the odor dilution bottle. Finally the odorized air is mixed with clean air, resulting in a new odor concentration in the airflow and incorporating new perturbations as a consequence of mixing the two airflows. **(B)** The baseline, and amplitude errors in sine-waves at different frequencies is not captured by a linear model (grey line), but it is captured by a NARX model (black line)

One approach to modeling a dynamical system is to develop an analytic framework that captures its physical characteristics. For example, one could use fluid mechanics to model the movement and interactions of the airflow within our system. However, this involves complex calculations for each setup, and would be highly sensitive to many parameters of the apparatus. Instead, we adopted a non-linear systems identification technique to model our olfactory setup: the *non-linear autoregressive with exogenous input* (*NARX*) model (***Bomberger and Seborg, 1998***). We trained the model with sine waves of different frequencies. The NARX model succeeded where linear models failed. For example, the NARX model was able to capture how baseline increases as frequency increases (Fig. 2C) and how amplitude decreases as frequency increases (Fig. 2D). A linear model, by contrast, failed to capture both the baseline-frequency and the amplitude-frequency relationships (Fig. 2C, D).

### Automated control of temporal odor waveforms

The PID controller in the MFC can only compensate for the disturbances that occur before or within the MFC but not after. We thus needed to compensate the non-linearity between the odorized airflow that leaves the MFC and the olfactory stimulus at the animal. To do this, we added a second feedback loop and a second PID controller. Two-degrees of freedom PID controllers (2-DOF-PID) are known to outperform PID controllers in robustness to perturbations while retaining set-point tracking accuracy (***Cominos and Munro, 2002***; ***Araki and Taguchi, 2003***). We implemented the 2-DOF-PID in an architecture where the set-point is the desired odor concentration waveform and the feedback signal is measured with a photoionization detector (Fig. 3A). The measured odor concentration at the output is subtracted from the desired stimulus odor concentration, and this difference or error signal is the input to the 2-DOF-PID. The 2-DOF-PID computes the input to the MFC that minimizes the odor concentration error signal.

**Figure 3.**
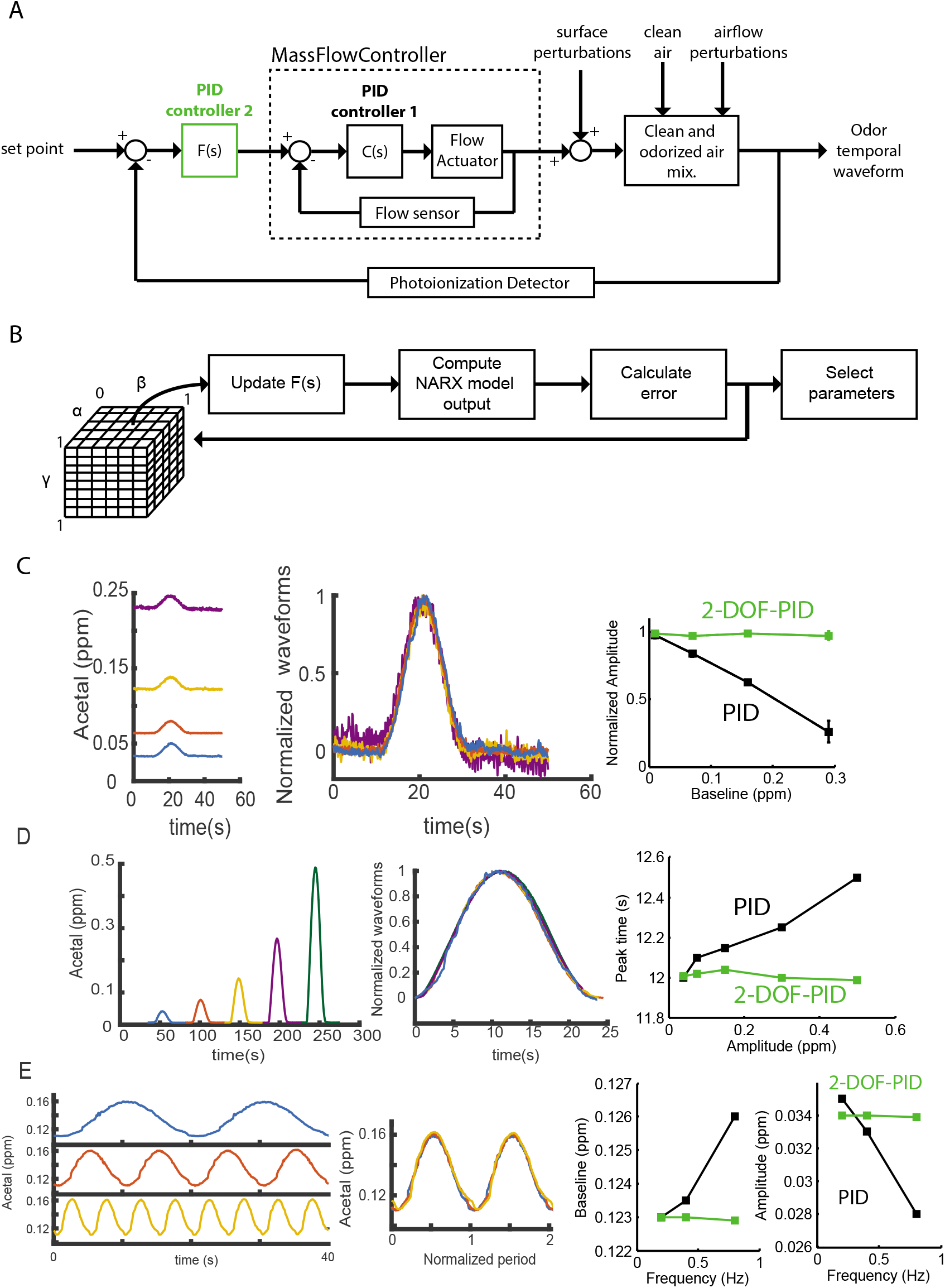
2-Degrees-of-freedom-PID odor control. **(A)** Implementing a second loop of PID control enables compensating for perturbations. **(B)** Recursive algorithm to use the NARX model to find sets of control parameters that compensate for perturbations. **(C)** Controlling identical odor waveforms with different baselines. Control error of 1-DOF-PID control (in black) vs. control error of 2-DOF-PID control (in green). **D** Controlling identical odor waveforms with different amplitudes. Control error of 1-DOF-PID control (in black) vs. control error of 2-DOF-PID control (in green). **(E)** Controlling identical sinewaves with different frequency. Control errors of 1-DOF-PID (in black), and of 2-DOF-PID (in green).

To calibrate the control parameters of the 2-DOF-PID, we use the NARX model to search for the three PID settings that result in the smallest error signals for a stimulus waveform (Fig. 3B). The parameters of the 2-DOF-PID are a function of the parameters of the PID controller that directly operates the MFC following equations described in (***Araki and Taguchi, 2003***). Once the parameters have been selected they are updated in the control software of the odor delivery system and tested experimentally. After the parameters have been tested, the system can work in open loop using the NARX model to simulate the photoionization detector feedback.

Our implementation of the 2-DOF-PID controller allowed us to define the amplitude, baseline, and frequency of the waveform of an olfactory stimulus. Unlike our attempts with a 1-DOF-PID controller, we were able to deliver identical odor waveforms with different baselines – superposition of the delivered waveforms after subtracting the baseline resulted in indistinguishable curves, and the amplitude of the delivered waveform was independent of baseline (Fig. 3C). We were also able to deliver odor waveforms with identical shape and fixed peak time but different amplitudes (green curve in Fig. 3D). Our earlier attempts to do this with a 1-DOF-PID controller resulted in different peak times (black curve in Fig. 3D). Finally, we were able to deliver sine waves of odor concentration with different frequencies and fixed baseline and amplitude (green curve in Fig. 3E). When we attempted to do this with a 1-DOF-PID controller, the baseline increased and the amplitude decreased with the stimulus frequency (black curve in Fig. 3E).

### Applications to the larval *Drosophila* early olfactory system

Being able to easily and precisely define an olfactory stimulus waveform in an airstream allows new analyses of neural processing and behavior. Here, we applied our setup to studies of the larval *Drosophila* olfactory system, which has the advantage of a small and highly accessible neural circuit. As in mammals, larval *Drosophila* olfactory receptor neurons (ORNs) typically express only one type of olfactory receptor (Or). Each of the 21 types of ORNs in the larva innervate an individual glomerulus in its antennal lobe (AL). To functionally image all 21 glomeruli we use transgenic animals with the calcium indicator GCaMP6m expressed under the control of the Orco-Gal4 driver (which labels all ORNs) and a spinning disk confocal microscope (Fig. 4A). We partially restrained the movements of a larva using an open-ended microfluidic channel, and positioned the head of the larva about 5 mm away from the outlet of the odor delivery apparatus (Fig.4-Supp.Fig. 1A).

**Figure 4.**
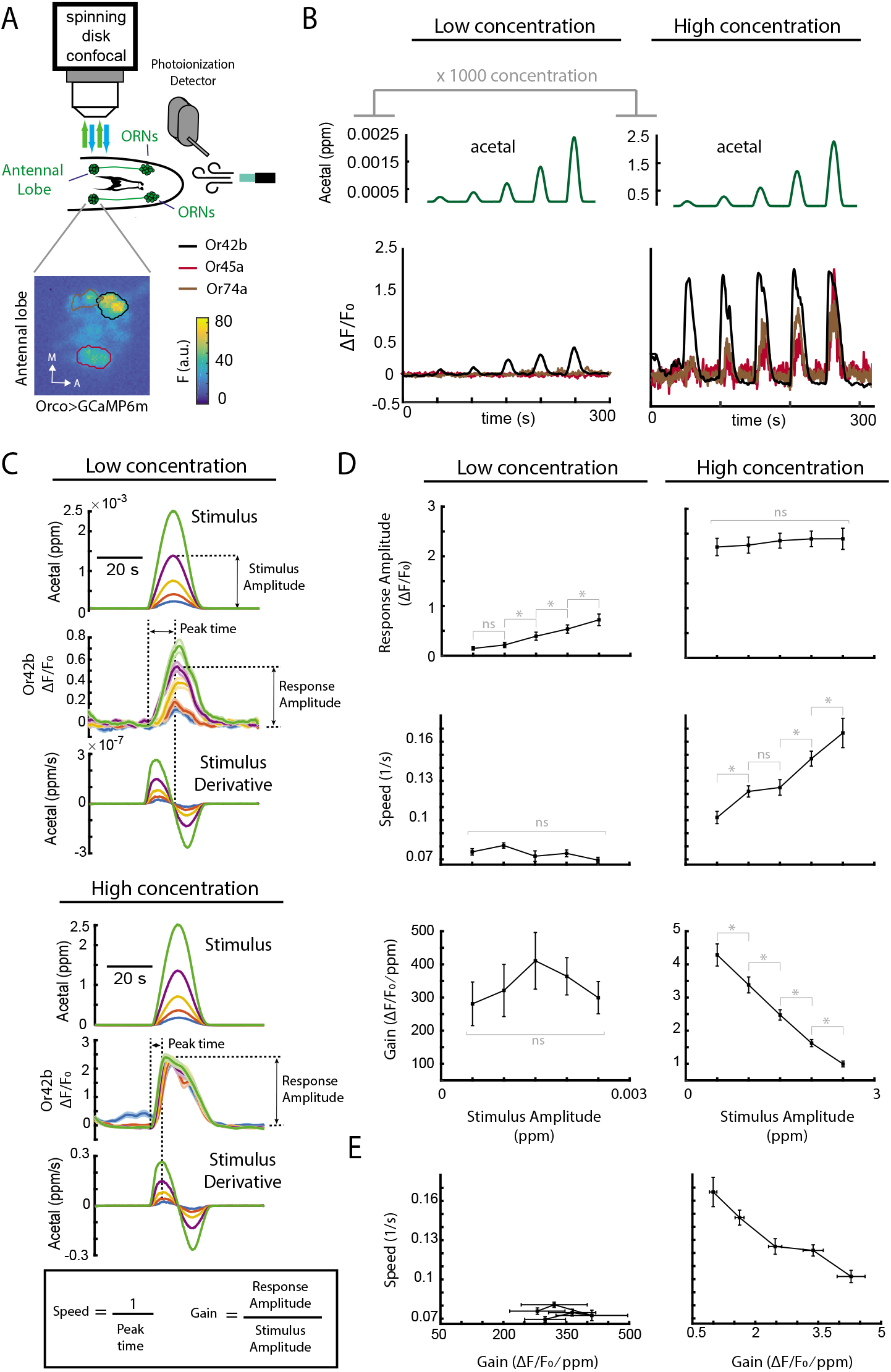
Dynamic adaptation at the ORNs. **(A)** Schematic of odor delivery system and microscopy system for larval *Drosophila* functional imaging in the Antennal Lobe (AL). **(B)** Responses of different olfactory glomeruli to odor sinusoidal ramps at low and high (1000x) concentrations. **(C)** Color coded odor stimuli, stimuli derivatives, and Or42b ORN calcium responses. Shaded regions in Or42b responses are the s.e.m. (n= 8 at low conc., n=12 at high conc., genotype: w^1118^, UAS-GCaMP6m, orco-Gal4) **D** Response, Amplitude, Speed and Gain vs stimulus amplitude, and Speed vs Gain. Error bars are the s.e.m. and indicate significantly different with p<0.05 in a Wilcoxon matched-pairs test. ‘n.s.’ indicates not significantly different. **Figure 4–Figure supplement 1.** Odor concentration and rate of change at the response peak. **(A)** Experimental preparation for *Drosophila* lavae. Larvae are embedded in a PDMS chip with the olfactory domes exposed and odorized arflow is delivered to the exposed head of the animal. **(B, C)** Odor concentration and derivative at the time point of maximum response amplitude. indicate significant difference with p>0.05 in Wilcoxon matched pairs test. ‘n.s.’ indicates not significantly different. Error bars are the s.e.m. **Figure 4–Figure supplement 2.** Dynamic Adaptation Model. **(A)** Dynamic Adaptation Model diagram. **(B)** Predicted response peak times match experimental results. **(C)** Predicted responses in red vs. experimental measurements in black.

#### Dynamic adaptation at the ORNs

ORNs can detect odor concentration changes over many orders of magnitude (***Wilson, 2013***). Most volatile compounds are specific to a few glomeruli at low concentrations and recruit additional glomeruli at higher concentrations (***Mathew et al., 2013***; ***Si et al., 2019***). Concentration-dependent recruitment of glomeruli is one mechanism for extending the range of stimulus detection (***Asahina et al., 2009***). Another mechanism, used for example in the retina, is dynamic adaptation (***Clark et al., 2013***). The neural responses of a dynamically adaptable sensor, such as rod or cone photoreceptor cells, are not only the result of integrating the stimulus but also the result of integrating the neural response itself. As a consequence of this integration, in dynamically adaptable sensors, gain suppression is accompanied by a speed-up of the neural response (***Clark et al., 2013***). This property enables dynamically adaptable sensors to encode information in their dynamics even when they are near saturation and they are no longer able to encode information in their response amplitude. Testing whether ORNs employ analogous mechanisms has been challenging without a way to interrogate their dynamics using identical stimuli differing only in amplitude.

In the *Drosophila* larva, acetal at low concentrations selectively activates the Or42b receptor (***Mathew et al., 2013***). We delivered acetal in two sets of odor concentration sinusoidal ramps during calcium imaging of ORNs in the glomeruli of the antennal lobe. The first set of stimuli was composed of low-concentration ramps that only activate the Or42b glomerulus. The second set was composed of high-concentration ramps that recruited two additional glomeruli (Fig. 4B). The low-concentration set of stimuli generated neural responses that increased linearly with the stimulus amplitude (Fig. 4C, D top panels). The neural response speed (defined as the reciprocal of the peak time) (Fig. 4C) and the gain (defined as the ratio between the response and stimulus amplitudes) were not significantly different among the responses to the low-concentration stimuli (Fig. 4D). In contrast, the neural responses to the high-concentration stimuli did not increase or decrease their response amplitude with the stimulus amplitude (Fig. 4C bottom panel, 4D top panel). Consistent with dynamic adaptation, we observed a speedup in the response that accompanied the suppression of the gain (Fig. 4D).

The trade-off exhibited by dynamically adaptable sensors can be visualized by plotting the gain versus the response speed. In Or42b ORNs, as in photoreceptors (***Clark et al., 2013***), gain suppression is accompanied by response speedup near saturation (Fig. 4E right). Below saturation, we observed no significant change in either gain or response speed (Fig. 4E left). The difference in response speed and gain is evident when comparing the responses at high and low concentrations: at low concentrations, the gain is hundred-fold higher than at high concentrations; at low concentrations, the response speed is half of the slowest response at high concentrations (Fig. 4E). At low concentrations, the peak response occurred near the maximum odor concentration when the stimulus time-derivative is near zero – this indicates an encoding of absolute odor concentration in ORN activity (Fig. 4C top). At high concentrations, the peak response occurred near the maximum odor concentration time-derivative – this indicates an encoding of the rate-of-change of odor concentration (Fig. 4C bottom).

The speedup in the ORN responses does not happen because a saturating odor concentration or odor concentration derivative is reached sooner when the stimulus amplitude is larger (Fig.4-Supp.Fig.1). We were also able to use a dynamic adaptation model similar to the one used for rods and cones (***Clark et al., 2013***) to fit the speed, gain, and dynamics of the Or42b-ORN neural responses to acetal (Fig.4-Supp.Fig.2 and Methods and Materials).

#### Odor contrast encoding

Unlike mammalian and adult *Drosophila* ORNs, larval *Drosophila* ORNs display incomplete adaptation. Even after several minutes of constant odor exposure, larval ORNs sustain an elevated firing rate (***Grillet et al., 2016***). Incomplete adaptation in sensors suggests a role of stimulus baseline in response dynamics (***Baylor and Hodgkin, 1974***). We investigated how ORN dynamics depend on the stimulus baseline.

We delivered acetal stimuli with different baselines but fixed amplitudes while recording calcium responses in the AL. At the lowest baseline, the calcium responses were significantly larger than at higher baselines (Fig. 5A). Both, the calcium response amplitude and the gain (defined as the ratio between response and stimulus amplitudes) decreased at higher stimulus baselines (Fig. 5B left) even though the stimulus amplitude was fixed (Fig. 5B center). With fixed amplitude and varying baseline, the stimulus contrast is variable. Using the definition of the Weber stimulus contrast used in vision studies (***Peli, 1990***), we calculate stimulus contrast as the ratio between the stimulus amplitude and the baseline (Fig. 5A). Higher stimulus contrast evoked stronger neural responses (Fig. 5B right).

**Figure 5.**
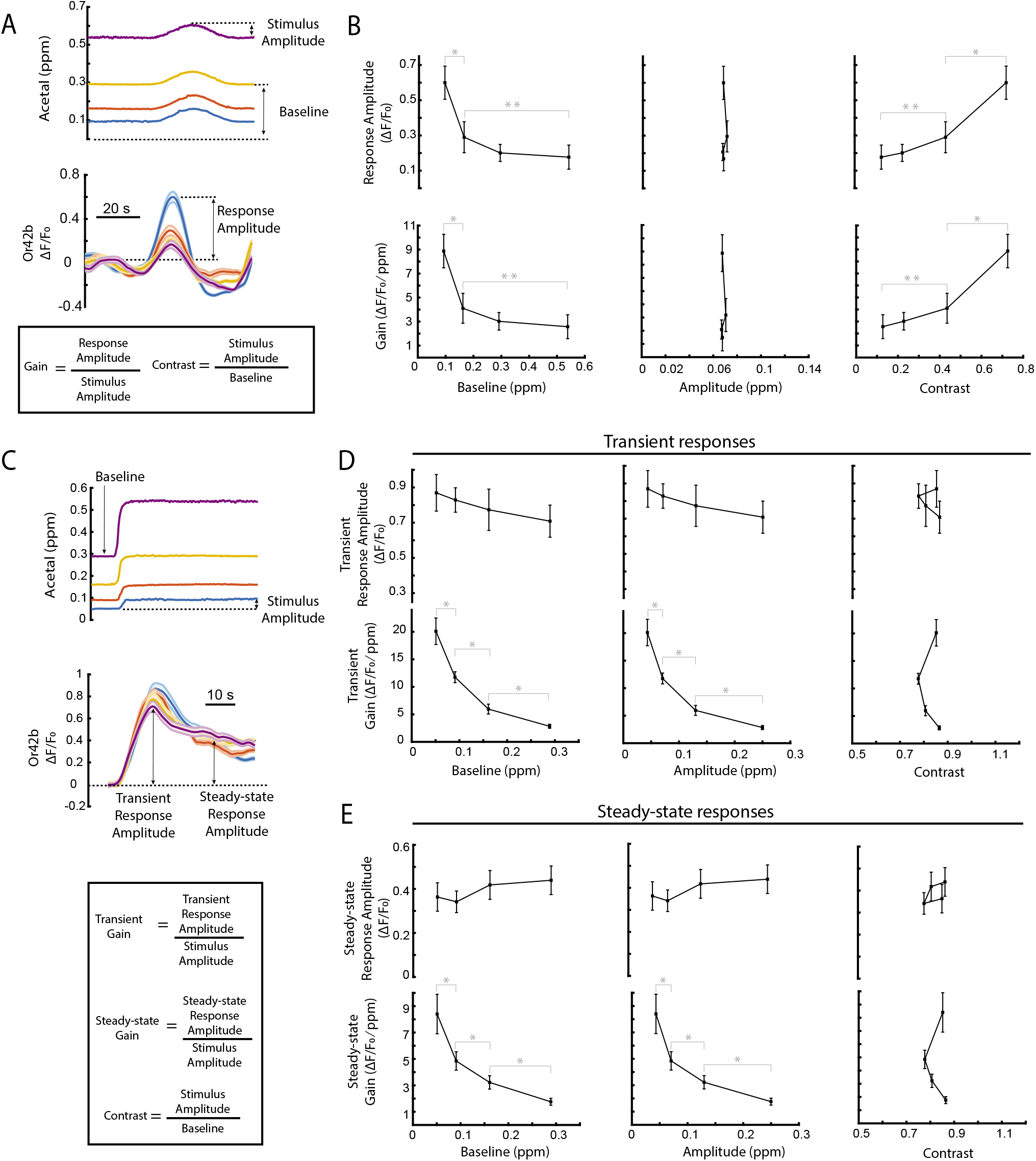
Odor contrast encoding. **(A)** Color-coded Or42b fluorescent responses to identical sinusoidal ramps with different baselines. (n=8 animals, genotype: w^1118^, UAS-GCaMP6m, orco-Gal4) **(B)** Response parameters with respect to stimulus baseline, amplitude, and contrast. **(C)** Color-coded Or42b responses to step stimuli with identical contrast at different baselines. (n=9 animals, genotype: w^1118^, UAS-GCaMP6m, orco-Gal4) **(D)** Response parameters with respect to stimulus baseline, amplitude, and contrast. In **(B)** and **(D)** error bars are the s.e.m., indicate significant difference with p<0.01 and ‘**’ indicate significant difference with p<0.05, calculated with Wilcoxon matched-pairs test. **Figure 5-Figure supplement 1.** Response peak time vs stimulus baseline. **(A)** Color-coded responses. **(B)** Response peak time vs. stimulus baseline. At lower baselines, responses are faster if amplitude is constant. **(C)** Color-coded responses. **(D)** Response peak time vs. stimulus baseline. If contrast is kept constant, response peak time is independent of stimulus baseline.

Next, we tested stimuli with different baseline but fixed stimulus contrast. Stimuli with fixed contrast evoked similar neural responses at different baselines (Fig. 5C). We separately analyzed the transient neural response (during and shortly after the odor step onset) and steady-state neural response (well after the odor step onset). In both regimes, the response amplitudes were independent of the stimulus baseline. The response-suppression that accompanies increases in stimulus baseline appears to be compensated by the response increase caused by the larger stimulus amplitude needed to maintain stimulus contrast (Fig. 5D, E). We observed a decrease in response gain with increasing baseline because, although response amplitudes stayed the same, the stimulus amplitudes needed to fix contrast increase with baseline.

Our results point to a model where larval *Drosophila* ORNs are specifically sensitive to odor contrasts in their environment independent of background odor concentration. The response to stimuli with fixed amplitude was also consistent with dynamic adaptation (Fig.5-Supp.Fig.1).

### Applications to larval *Drosophila* odor-guided navigation

In the study of odor-guided navigation, it has been difficult to deliver sensory input to behaving animals with the flexibility and precision needed to probe the dependence of behavioral dynamics on stimulus dynamics. One approach is to model the physics of the odor gradient, and estimate the sensory experience of the animal navigating within it. For example, stimulus dynamics can be estimated using the diffusion of an odor from a point source while monitoring an animal’s behavior in a closed environment (***Gomez-Marin et al., 2011***). Optogenetics circumvents this challenge by artificially activating specific neurons while recording behavior (***Schulze et al., 2015***; ***Hernandez-Nunez et al., 2015***; ***Gepner et al., 2015***). However, optogenetics does not necessarily evoke the neural dynamics that occur in response to environmental odors.

In a previous study, ***Gershow et al*.** (***2012***) engineered a behavioral apparatus that is able to generate spatial linear gradients with air currents that are small enough to avoid rheotactic responses in larval *Drosophila*. Here, we applied our control method to the behavioral chamber from ***Gershow et al. (2012)*** to precisely manipulate temporal odor dynamics while recording freely moving larval *Drosophila*.

#### Controlling odor waveforms in a behavioral arena

We adapted our system in Fig. 1 for the behavioral arena developed by ***Gershow et al*.** (***2012***). The only hardware difference before odor delivery is the substitution of the Y junction of Fig. 1A for a mixing chamber (Fig. 6A). Two massflow controllers are used to deliver odorized and clean air flows into the metallic mixing chamber before introduction to the behavioral arena. The laminar odorized air flow travels through the behavioral arena before exiting through an outlet. At the outlet, a photoionization detector measures odor concentration (Fig. 6A). We also measured odor concentration at several points within the arena to calibrate the controller and to account for time delay as a temporal waveform of odor concentration travels through the behavioral arena (Fig.6-Supp.Fig.1 and Methods and Materials). Using our 2-DOF-PID control strategy (see above) we were able to deliver odor stimuli 20 times faster than the stimuli generated with traditional PID control (Fig.6-Supp.Fig.1).

**Figure 6.**
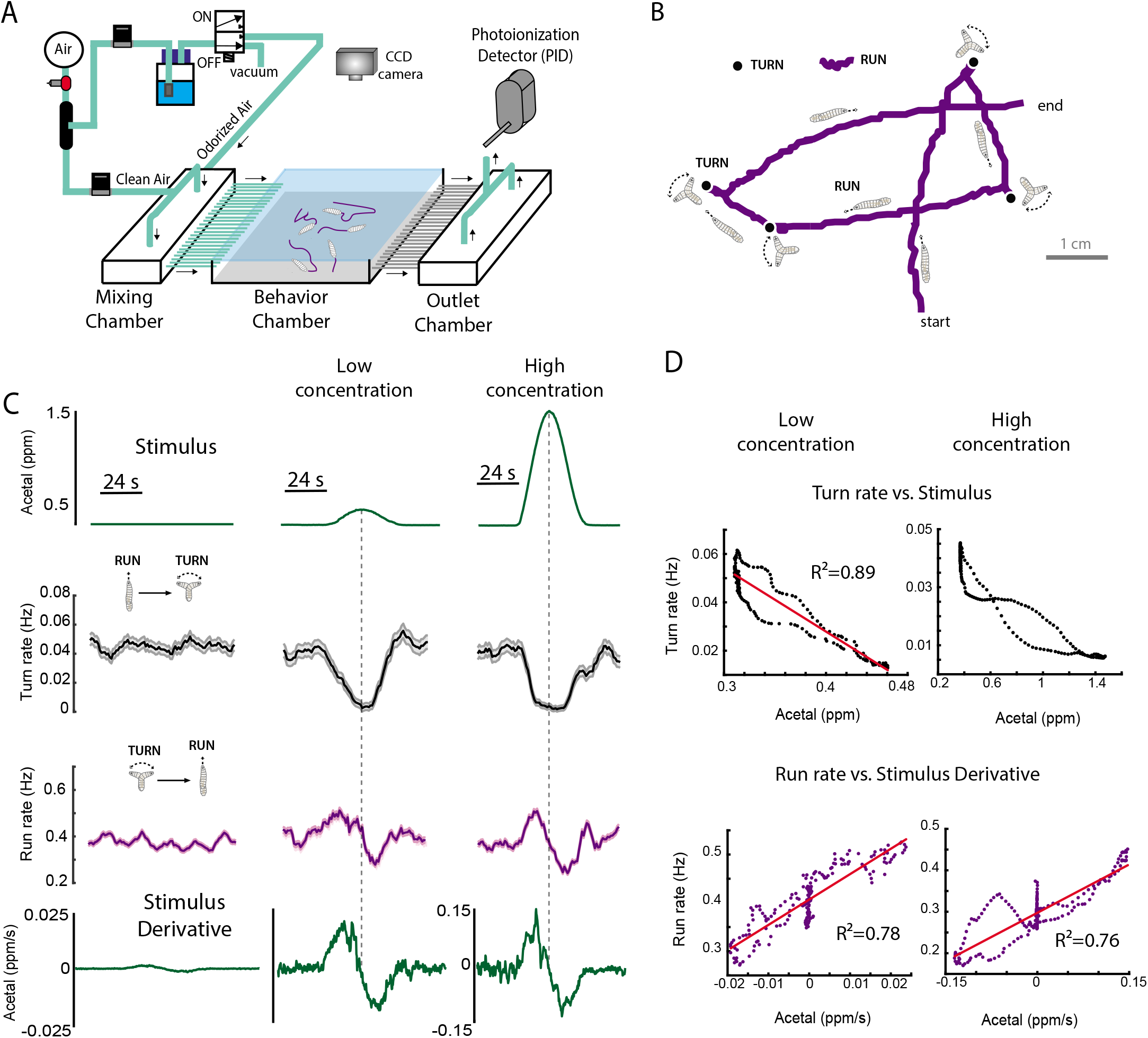
Controlling odor waveforms in a behavioral chamber. **(A)** Schematic representation of the odor delivery apparatus and behavioral chamber. **(B)** Larval *Drosophila* navigate their environment by alternating between two motor programs: runs (in purple) and turns (black dots). **(C)** Behavioral responses to low and high concentration sinusoidal odor ramps. (n=95 at low conc., n=81 at high conc., Genotype w^1118^) **(D)** Correlation between behavioral dynamics and the stimulus or the stimulus temporal derivative. Pearson correlation coefficients are shown in each panel. **Figure 6–Figure supplement 1.** Odor waveforms in a behavioral chamber. **(A)** At different distances of the delivery valves, identical odor waveforms have increasing latencies. **B** 2-DOF-PID controllers enable the delivery of faster odor waveforms than 1-DOF-PID controllers.

In spatial odor gradients, olfactory preference is inferred by the direction of movement and aggregation of behaving animals (***Kreher et al., 2005***, ***2008***; ***Vogt et al., 2021***). In temporal odor gradients, olfactory behavioral responses must be inferred by animal movements in response to increasing and decreasing odor concentrations over time (***Hernandez-Nunez et al., 2015***; ***Gepner et al., 2015***). Larval *Drosophila* olfactory navigation involves the regulation of two motor states, runs during which larvae crawl forward in peristaltic movement and turns during which larvae stop forward movement and sweep their heads until they select a new run direction (***Gershow et al., 2012***; ***Luo et al., 2010***; ***Kane et al., 2013***) (Fig. 6B). When larvae encounter increasing odor concentrations of an attractant during a run (akin to moving up a spatial gradient of the odor), they lower the likelihood of starting a turn, thereby lengthening runs in favorable directions. When larvae encounter increasing odor concentrations of an attractant during a turn, they increase the likelihood of starting a new run, thereby starting more runs in favorable directions (***Hernandez-Nunez et al., 2015***; ***Gepner et al., 2015***). Thus, a temporal increase in the concentration of an attractant should both suppress transitions from runs to turns and stimulate transitions from turns to runs. A temporal decrease in attractant concentration should have the opposite effects.

To study how odor stimuli dynamics map to behavioral dynamics, we conducted experiments with ~ 20 larvae that freely crawled within the behavioral arena while sinusoidal ramps of odor concentration were delivered. For control experiments, where we wanted a non-odorized airflow, we put clean water in the odor dilution bottle. This created small changes in humidity, but these changes were not large enough to trigger measurable behavioral responses (Fig. 6C right). A low concentration sinusoidal stimulus of acetal triggered a decrease in the rate of run-to-turn transitions (turn rate) and an increase in the rate of turn-to-run transitions (run rate) during the rising phase of the stimulus (Fig. 6C center), consistent with previous studies that report acetal as an attractant (***Mathew et al., 2013***). During the falling phase, we observed an increase in the turn rate and a decrease in the run rate (Fig. 6C center). Behavioral responses to sinusoidal stimuli at high concentration resulted in similar trends (Fig. 6C left).

At low concentrations, turn rate peaked near the maximum odor concentration of the stimulus waveform (Fig. 6C center). In this regime, turn rate is sensitive to the absolute value of odor concentration, consistent with the calcium response exhibited by the Or42b-ORN at low concentrations (Fig. 4C top). Turn rate is highly correlated with stimulus dynamics (Fig. 6D). At high concentrations, turn rate peaked near the maximum derivative of odor concentration (Fig. 6C right). In this regime, turn rate is sensitive to the rate of change of odor concentration, consistent with the calcium responses exhibited by Or42b-ORN at high concentrations (Fig. 4C bottom). At high concentrations, the shape of the turn rate response is closer to a saturated version of the stimulus and so stimulus and response are not linearly correlated (Fig. 6D). The run rate is correlated with the time-derivative of the stimulus at both low and high concentrations (Fig. 6D bottom).

#### Mapping neural dynamics to behavioral dynamics

The precise control of olfactory stimuli enables the delivery of identical odor waveforms for neurophysiology and behavior, facilitating the study of sensorimotor transformations by comparing results from different experiments. We measured neural responses in larval antennal lobe (AL) neurons to determine whether behavioral dynamics that are not represented by sensory neuron dynamics might be accounted for by the dynamics of second-order neurons.

The wiring diagram of the larval AL neurons has been mapped with synaptic resolution (***Berck et al., 2016***). The AL contains two major input-output pathways. One pathway is composed of glutamatergic picky local neurons (pLNs) that receive inputs from a few glomeruli and modulate multiglomerular projection neurons (mPNs). The other pathway is composed of GABAergic broad local neurons (bLNs) that receive inputs from all glomeruli and modulate uniglomerular projections neurons (uPNs). The uPN pathway has been shown to mediate behavioral responses to attractants (***Vogt et al., 2021***) (Fig. 7A). We asked how the neural dynamics of the uPN pathway might be related to behavioral dynamics in response to an attractant.

**Figure 7.**
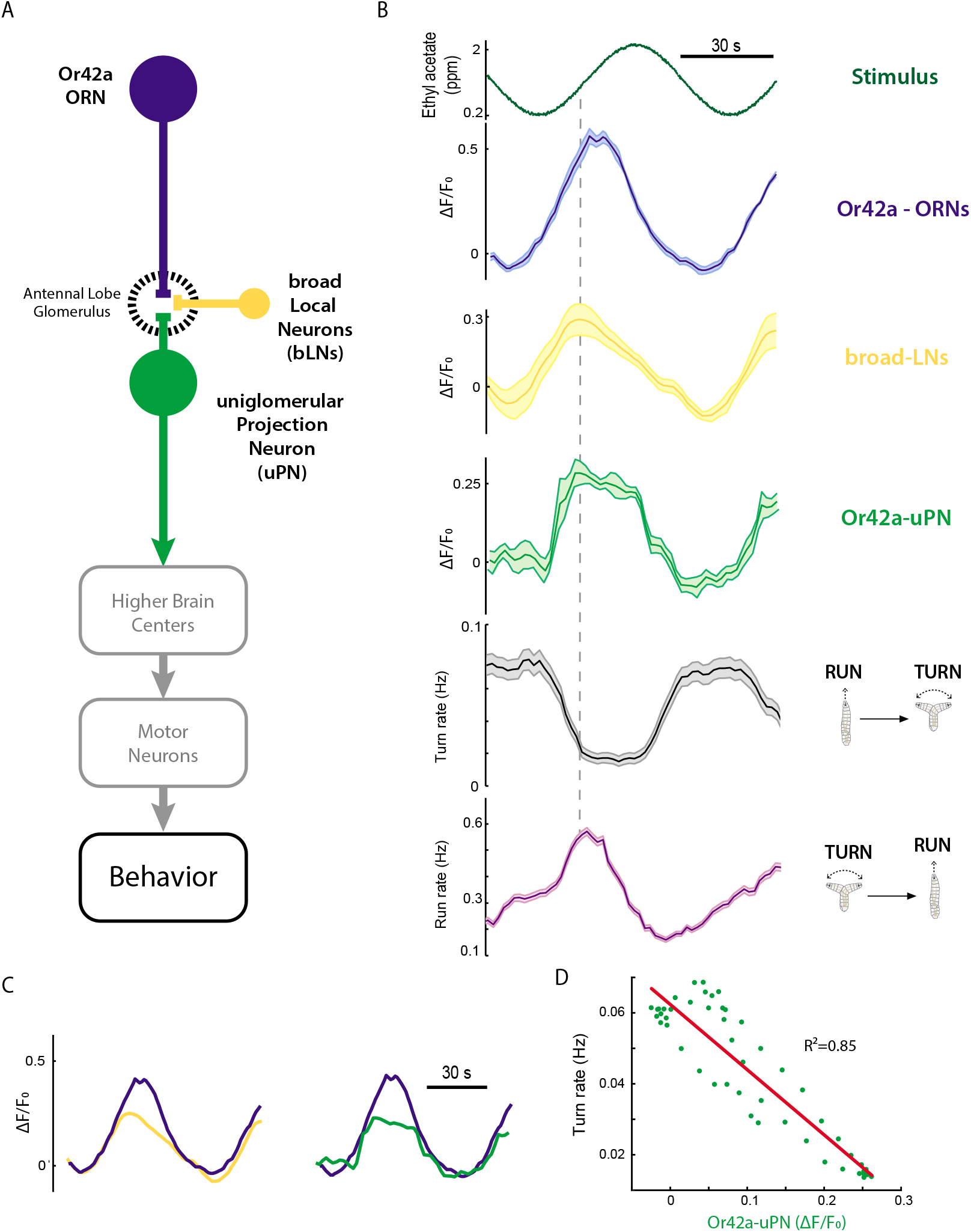
Neural dynamics and behavioral dynamics. **(A)** Schematic representation of the uniglomerular olfactory circuit in larval *Drosophila*. **(B)** Calcium responses in ORNs, broad LNs, uPNs, and behavioral resposnes of wildtype larvae. (Genotypes: (ORNs) w^1118^;Or42a-Gal4/orco-RFP;UAS-GCaMP6m (broad-LNs) w^1118^;R51E09-Gal4/orco-RFP; UAS-GCamP6m/+ (Or42a-uPNs) w^1118^;GH146-Gal4;UAS-GCaMP6m and w^1118^ wildtype for behavior) (n=8-12 for calcium imaging, n=97 for behavior). **(C)** Overlaid mean ORN, broad-LN, and uPN responses. **(D)** Linear correlation between the Or42a-uPN response and the turn rate. **Figure 7-Figure supplement 1.** Identification of uniglomerular circuit components. **(A)** Antibody staining with Alexa 488 anti-GFP, Alexa 555 anti-GABA in R51E09-Gal4; UAS-GFP first instar larvae. nc82 was used as a reference. **(B)** Light microscopy vs electron microscopy reconstruction of the uPNs. (GH146-Gal4/LexAop-RFP;R12E10-LexA/UAS-GFP)

**Figure 8.**
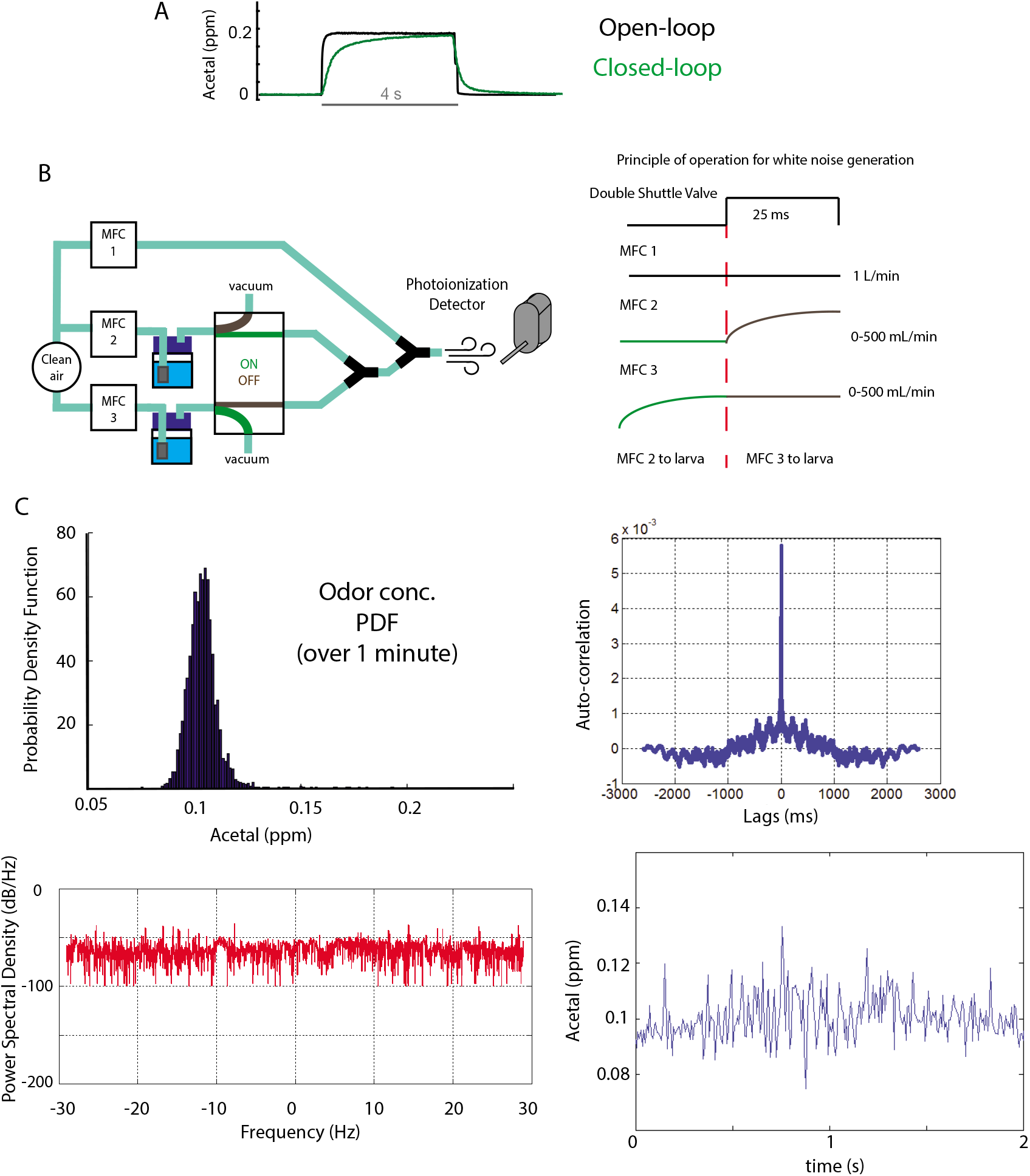
Odor delivery speed. **(A)** Closed- vs. open-loop delivery speed for step stimuli. **(B)** Olfactometer configuration for odor concentration white noise process stimuli. **(C)** White process distribution, autocorrelation time, power spectrum, and temporal resolution.

We compared neural and behavioral responses to identical sinusoidal waves of ethyl acetate, a widely used attractant with a fruity smell (***Kreher et al., 2008***; ***Fishilevich et al., 2005***). Ethyl acetate principally activates ORNs that express Or42a but also recruits other ORNs (***Kreher et al., 2008***; ***Si et al., 2019***).

To measure neural responses, we expressed GCaMP6m under the control of Gal4 drivers in the Or42a ORN (Or42a-Gal4), the bLNs (R59E09-Gal4), and the uPNs (GH146-Gal4). The identity of labeled neurons was easily matched to the AL connectome based on neurite morphology and antibody stainings (Fig.7-Supp.Fig.1). The neural responses of the Or42a ORNs, the bLNs, and the uPNs peaked during the rising phase of the sinusoidal odor waveform and reached a minimum during the falling phase (Fig. 7B). The bLNs and uPNs peaked before the ORNs (Fig. 7C). Thus, these second-order neurons encode derivative components of the ORN responses. While the ORN and bLN responses were smooth, the uPN responses resemble a rectified version of the ORN response.

We measured behavioral responses to the same sinusoidal odor waveform. We found that the turn rate response decreases during the rising phase and increases during the falling phase. In contrast, the run rate response increases during the rising phase and decreases during the falling phase. These responses are consistent with the general behavioral response to an attractant (Fig. 7B). Or42a-ORN response dynamics are not directly reflected in turn and run rate dynamics. Run rate coincides in peak time with bLNs and uPNs (dashed line in Fig. 7B); following the peak, we note that the run rate decays faster than the neural responses. The dynamics exhibited by turn rate are highly correlated with uPN neural responses (Fig. 7D), suggesting that the transformation needed to shape ORN dynamics into turn rate dynamics has occured in the AL, the first olfactory processing center in larval *Drosophila*.

## Discussion

To understand how the brain transforms environmental cues into behavioral dynamics, it is crucial to be able to precisely control the temporal dynamics of sensory stimuli. The study of vision, audition, and other sensory modalities has benefited from experimental setups allowing precise temporal control of sensory stimuli to probe how circuits shape behavior (***Clemens et al., 2018***; ***Badwan et al., 2019***; ***Klein et al., 2015***; ***Hernandez-Nunez et al., 2020***). The technical difficulty of controlling odor temporal dynamics for neurophysiology and behavior experiments has drastically constrained the study of the temporal structure of olfactory sensorimotor transformations. Here we leveraged control engineering and systems identification techniques to precisely control temporal odor waveforms. We then show the experiments this advancement enables by studying the larval *Drosophila* olfactory system and navigational behavior.

### Technological advancement

Since the discovery of the olfactory molecular receptors in rodents and flies (***Buck and Axel, 1991***; ***Vosshall et al., 1999***; ***Clyne et al., 1999***), the tools used to study olfactory information processing have grown in sophistication. In the first two decades post-receptors discovery (until 2011), the field of *Drosophila* olfaction relied mainly on natural diffusion of an odor source in a closed environment to study olfactory behaviors (***Fishilevich et al., 2005***; ***Kreher et al., 2008***) and on solenoid valves to deliver odor puffs to study sensory encoding (***Asahina et al., 2009***; ***Kreher et al., 2005***). Those techniques helped us learn plenty about the neural and molecular mechanisms underlying odor preference, and about combinatorial encoding of odor identity; however they were less effective in helping us understand the temporal processing of olfactory stimuli.

In the last decade, many studies have developed new ways of controlling odor dynamics to study the temporal structure of neural processing and sensorimotor transformations in *Drosophila*. ***Nagel and Wilson*** (***2011***) used linear filters to model an odor delivery system and more reliably deliver square odor pulses to *Drosophila* while using electrophysiology to record sensory neuron activity. More recently, ***Álvarez-Salvado et al*.** (***2018***), used proportional valves to generate ramps and turbulent odor plumes while recording *Drosophila* behavior. ***Schulze et al***. (***2015***) and ***Gorur-Shandilya et al*.** (***2017***), used massflow controllers to indirectly control odor concentration while measuring sensory neuron activity. These approaches moved the field from the use of odor puffs to the use of temporal odor waveforms. However, a method to precisely define a desired temporal waveform has been lacking. Precise delivery of a defined temporal waveform would facilitate comparisons between experiments, such as comparing neural responses in a physiology experiment with motor responses in a behavioral experiment.

We have developed a control system for odor waveforms that enables us to independently control multiple salient variables in olfactory computation including the baseline, amplitude, and temporal structure of odor concentration waveforms. Independent control of these parameters enables clean dissection of odor temporal processing capabilities in olfactory neural circuits. The flexibility of our method for both neurophysiology and behavioral studies, enables delivering identical odor stimuli in different experiments which is needed to rigorously establish correlations between odor, neural, and behavioral dynamics.

### Technical limitations

Most control systems, like PID controllers, have a trade-off between set-point tracking and robustness to perturbations (***Araki and Taguchi, 2003***). Control systems that are better at set-point tracking will under-perform when responding to perturbations. When delivering odors, there are many sources of perturbation because of the inherent properties of airflow dynamics. PID controllers are particularly susceptible to the trade-off caused by the non-linear transformation from odorized airflow to final odor concentration in odor delivery systems. We have found that a 2-DOF-PID controller alleviates the performance-robustness trade-off in odor delivery.

The 2-DOF-PID controller alleviates the compromise between set-point tracking and robustness to perturbations, but still has constraints. Set-point functions that are not continuous are more likely to create set-point tracking delays. This limits the speed of presentation of odor waveforms, but does not limit the capacity of the apparatus to deliver fast odor fluctuations in open loop. In open loop, we are able to deliver white noise odor stimuli at 40Hz (Extended Methods).

### Temporal processing of olfactory information

We have shown how precisely controlled odor waveforms can be used to investigate the temporal encoding of olfactory stimuli. Using the larval *Drosophila* Or42b ORN, we show that far from saturation (at low odor concentrations) the amplitude of the olfactory stimulus modulates the amplitude of the neural response but not the dynamics. Near saturation, the amplitude of the stimulus modulates the dynamics of the response but not the amplitude. Previous studies in larval *Drosophila* found invariant dynamics in ORN odor encoding, consistent with our findings far from saturation (***Si et al., 2019***).

Near saturation, gain suppression comes accompanied by a speed-up of the response, consistent with a phenomenon called the “gain-bandwidth trade-off”, a property of dynamically adaptable sensors (***Clark et al., 2013***). Second order olfactory neurons in *Drosophila* are known to encode a component of the time-derivative of ORN activity (***Kim et al., 2015***). The non-linear adaptation of Or42b-ORNs near saturation may be a mechanism of transmitting derivative information when the sensor response amplitude is saturated.

We have also shown that larval ORNs directly encode stimulus contrast. Consistent with findings in adult *Drosophila* (***Gorur-Shandilya et al., 2017***), larval ORNs are less responsive to odor concentration changes at higher odor baselines. The precise drop-off of sensitivity with baseline enables the Or42b ORNs to encode contrast: stimuli with the same contrast but different baselines produced indistinguishable neural responses. This finding underscores the previously reported Weber-Fechner law in ORN encoding (***Gorur-Shandilya et al., 2017***). Flies may need to suppress the gain of ORNs at higher baselines to be better tuned for contrast detection.

### Olfactory sensorimotor transformations

Olfactory behavior has long been studied using fixed odor sources and measuring animal aggregation near or away from the odor source (***Asahina et al., 2009***; ***Kreher et al., 2005***; ***Mathew et al., 2013***; ***Vogt et al., 2021***). Aggregation assays provide a scalar metric of preference but not information about behavioral dynamics. We have developed a control technique for odor stimuli that allows us to precisely probe the temporal characteristics of behavioral and neural dynamics which is needed to illuminate computation in olfactory circuits. Olfactory behavior in the *Drosophila* larva is straightforward to quantify as transitions between two motor programs, runs and turns. We have determined how run-to-turn transitions (turn rate) and turn-to-run transitions (run rate) depend on the temporal properties of an odor waveform. Two odor waveforms of identical shape but different amplitude revealed that, at low concentrations, the dynamics of the turn rate follow the odor waveform – this is consistent with ORN responses at low odor concentrations. At high odor concentrations, turn rate peaks near the maximum derivative of the odor waveform – this pattern is consistent with ORN responses at high odor concentrations. Run rate dynamics at both high and low odor concentrations are correlated with the time-derivative of the odor waveform.

To understand how animal brains transform odor stimuli into behavioral responses, we need to locate neural dynamics relevant for behavior in the neural circuits that process olfactory information. For the sinusoidal stimulus we used, ORN response dynamics are not directly correlated with run or turn rate. We found that the dynamics of turn rate are highly correlated with the activity of uniglomerular projection neuron (uPN). Run rate peaked simultaneously with broad local neurons (bLNs) and uPNs, but their decay dynamics were faster than those of all the neural responses we measured. Run rate may be computed elsewhere in the neural circuit.

The stochastic nature of olfactory behaviors makes it often necessary to record from dozens or even hundreds (as in the case of larval *Drosophila*) of animals to estimate behavioral responses to a stimulus. A key advantage of our method is the high reproducibility of the stimulus waveform, making it easier to draw comparisons between neurophysiology experiments with a small number of animals (n=8-12) and behavior experiments with large numbers of animals (n=80-120).

### Significance and future directions

Precise stimulus control that spans physiology and behavior is needed to dissect neural circuits for sensorimotor transformations. The techniques for odor delivery that we have introduced here enable using identical odor waveforms for functional imaging and behavior experiments. We have also shown how systematic variation of stimulus waveform parameters, such as baseline, amplitude, or contrast, without altering other stimulus features enables dissecting the dynamics of sensory neurons and behavior. Combining this approach with the wiring diagrams and genetic toolboxes available in *Caenorhabditis elegans*, or larval and adult *Drosophila melanogaster* can illuminate how neuromodulators and interneurons sculpt the circuit dynamics that give rise to navigational decisions and behavioral states.

In nature, multiple odors vary dynamically in time and space. Future efforts to augment the performance of odor delivery devices should address the independent control of multiple simultaneous odor waveforms. This will enable the simultaneous study of the combinatorial and temporal integration properties of olfactory systems.

## Methods and Materials

### Control algorithm and parameter tuning

The control algorithm includes two feedback loops that control two separate variables, flow and odor concentration. Both controllers implement, proportional, integrative, and derivative (PID) action, and use the error signal (the difference between the desired and the measured quantity) as input. While the PID controller in the MFCs only act locally, the PID for odor control uses the the odor concentration at the animal location as input. The delay between changes in the input to the MFC and the output of the system constitute the timescale bottleneck of the system.

The parameters of each controller can be tuned independently, following set-point tracking and overshoot and oscillation criteria as described in (***Zhong, 2006***). However, we found it much simpler to use the 2-DOF-PID feedforward compensator tuning equation described in (***Araki and Taguchi, 2003***), where the parameters of the PID controller 2 depend on the parameters of PID controller 1. The control equations of each controller are as follows:

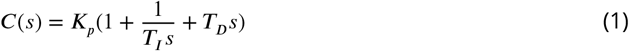

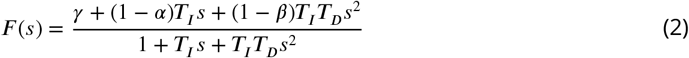

In equation 1, *C*(*s*) is the transfer function of PID controller 1. The Laplace transform was used to transform the function from the time real variable *t*(time) to the complex variable *s* (frequency). The constants *K_P_*(Proportional component), *T_I_*(Integral component), and *T_D_*(Derivative controller), are manually tuned to eliminate overshooting of the system and guarantee convergence between the desired set-point and the experimental output. PID-contorller 2 parameters relate to the parameters of PID controller 1 through equation 2. The transfer function (*F*(*s*)) depends on the PID parameters of controller 1 and on the new parameters *γ*, *α*, and *β*.

The new parameters *γ*, *α*, and *β* are tested recursively in a simulation using a NARX model of the olfactometer.The three parameters are sampled from 0 to 1 and we run simulations with thousands of different parameter combinations until the result fits the desired signal with *R*^2^ > 0.9. Then we use these parameters experimentally to verify the accuracy of the tracking.

### Systems identification techniques

In order to optimize control parameters we needed a computational model that captures the errors of the system. We used the Matlab Systems Identification toolbox to test linear transfer functions, ARX (auro-regresive with exogenous input) models, and NARX (non-linear ARX) models. The NARX model captures these erros and therefore can be used to optimize the controllers computationally. The custom MATLAB code and results can be found in https://github.com/LuisM6/OdorTemporalControl2021.

### Running the system in open-loop

Using the NARX model to calculate the output using the input history we can run the system without the photoionization detector feedback with negligible loss in accuracy. The Virtual Instruments needed to run the setup can also be found in the LabVIEW folder in https://github.com/LuisM6/OdorTemporalControl2021.

### Neurophysiology experiments

Calcium imaging experiments were conducted using a spinning-disk confocal microscope (Nikon microscope and ANDOR spinning disk) and a 60x 1.2-N.A water immersion objective (Nikon Instruments LV100; Andor). Early first-instar larvae were injected in a channel of a PDMS chip using a syringe such that they were immobilized by the channel walls, while leaving their olfactory domes exposed outside of the channel (Fig. 4-Figure supplement 1). The bottom of the PDMS chip was fixed on a metallic stage and a stainless steel tubing carrying the odorized airflow was positioned at 5 mm from the larval olfactory domes and with the axis of the larvae aligned with the stainless steel tubing. The odor control software was synchronized with the volumetric image acquisition using National Instruments Data Acquisition Cards (NIDAQs). The software is in the the github page of this manuscript (https://github.com/LuisM6/OdorTemporalControl2021) and can be used as described in the README file.

### Behavior experiments

The odor delivery system used for behavior is identical to the one described for physiology but with the output connected to the hermetic behavioral arena previously described in (***Gershow et al., 2012***). In this apparatus, instead of mixing the odorized and clean airflows using a Y junction, the clean air enters a custom built metallic chamber and the odorized airflow enters a different chamber and then passes through an array of 29 valves to mix with the clean air and be released in the behavioral arena. The photoionization detector was used with a metallic grid that can be clamped to the behavioral arena while keeping it hermetic and enables the insertion of the photionization detector probe at different locations of the arena (see (***Gershow et al., 2012***).

For each behavior experiment, second instar larvae were selected based on spiracle development under a dissection microscope. Larvae were then washed twice before transferring them to a thin (1.5mm) 2 percent agar, and 0.2 percent activated charcoal layer previously deposited inside the behavioral chamber. Larvae movements were recorded using a high-pixel-density CCD camera (Mightex) at 4Hz while illuminated with red light with LED bars located on the sides (dark field illumination). The behavioral recordings and stimuli were generated and synchronized using custom LabVIEW software (https://github.com/LuisM6/OdorTemporalControl2021).

### Calibration of photoionization detectors

The Aurora miniPID-200b sensor was calibrated for different signal levels using a set of 3 precalibrated piD-TECH eVx photionization sensors that were tuned for detection at different baseline levels. The detection limits of each eVx sensor were 10,1.5, and 0.5 parts per billion respectively, and their response times 4 seconds. The miniPID-200b was able to detect odor concentrations accurately in all regimes once calibrated, with a response time near 10 miliseconds.

### Neurophysiology data analysis

To analyze the calcium imaging data first we conducted motion correction using moco-master (***Dubbs et al., 2016***) to computationally cancel small movements of larvae within the PDMS chip (https://github.com/LuisM6/OdorTemporalControl2021). The motion-corrected data was then segmented and quantified using constrained non-negative matrix factorization (CNMF) (***Pnevmatikakis et al., 2016***) (https://github.com/LuisM6/OdorTemporalControl2021).

### Behavior data analysis

Behavioral data was segmented, and larval contours and position over time were extracted using MAGAT Analyzer (***Gershow et al., 2012***) (https://github.com/samuellab/MAGATAnalyzer). The quantification of run and turn events was then implemented with custom MATLAB code (https://github.com/LuisM6/OdorTemporalControl2021). Only a 10×10 cm square at the center of the arena was used for calculating behavioral responses, as odor dynamics were less reliable near the edges in our measurements.

### Dynamic Adaptation Model

We implemented a dynamic adaptation model following (***Clark et al., 2013***). The DA model equation is as follows:

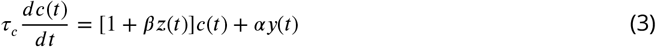

In this equation, *τ_c_*, *β*, and *α* are positive constants. The calcium response *c*(*t*) depends on *y*(*t*) and *z*(*t*) that depend on the temporal odor concentration *s*(*t*) as follows:

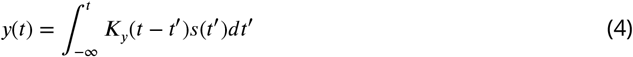

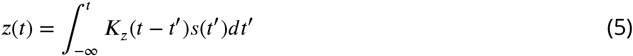

The kernels *K_y_*(*t*) and *K_z_*(*t*) integrate to unity and are the product of exponential and monomial functions as indicated in the methods of (***Clark et al., 2013***).

### Fly husbandry and genetics

Flies were raised at 50% humidity and constant temperature (22 °C), on standard cornmeal agar-based medium. To conduct crosses and collect larvae, adult *Drosophila melanogaster* were transferred to collection cages (Genesee Scientific). One side of the cage held a grape juice agar plate and fresh yeast paste. Flies laid eggs on the grape juice plate for 2 days. Then the plate was removed to collect larvae. For calcium imaging experiments first instar larvae were selected, and for behavior experiments early second instar larvae were selected based on spiracle development.

The genotypes of fly stocks used in this study:

**Effectors:** UAS-GCaMP6m in the second chromosome: w[1118]; P[y[+t7.7] w[+mC]=20XUAS-IVS-GCaMP6m]attP40 (from BDSC 42748). UAS-GCaMP6m in the third chromosome: w[1118]; PBac[y[+mDint2] w[+mC]=20XUAS-IVS-GCaMP6m] VK00005(from BDSC42750). UAS-GCaMP6m in orco-RFP background: UAS-mCD8::GFP; Orco::RFP (from BL63045).
**Gal4-Drivers:** w[1118];Or42a-Gal4(BL9971). w[1118];Orco-Gal4(BL23292).y[1] w[1118]; GH146-Gal4 (BL30026). w[1118];R51E09-Gal4 (JFRC).

## Acknowledgments

We want to thank Christopher Tabone, Guangwei Si, ans Matthew Berck for feedback on the project, and helpful discussions. We also want to thank Christopher Tabone for the staining of broad LNs. We thank Rachel Wilson, Sandeep Robert Datta, Alex Schier, and Andrew Murray for helpful discussions and advice during the development of the project. We thank Katrin Vogt, David Zimmerman, Albert Lin, and Jessleen Kanwal for comments on preliminary versions of the manuscript. We acknowledge the Bloomington *Drosophila* Stock Center (NIH P40OD018537). L.H-N. and A.D.T.S. were supported by the NIH R01 GM130842-01 grant.

## Contributions

L.H-N. and A.D.T.S. designed the study and wrote the manuscript. L.H-N. interpreted the results, developed the experimental techniques, wrote the software, analyzed data, and conducted the experiments.

## Extended Methods

The design of an olfactometer requires solving intermediate technical problems. Here, we explain the pressure compensation problems that typically occur with olfactometers and the solutions we devised, next we explain the contamination problems the system may experience, then the speed bottlenecks for set-point tracking, and finally how to generate faster signals in open loop.

### Pressure compensation

When using or developing an olfactometer, pinch solenoid valves are often the choice for controlling the delivery of an odor puff. Using those valves results in building up pressure in the tubings of the system and having bursts of odor concentration after valve openings. In addition, when pinch valves are open and clean air is flowing while the odorized air branch is set to zero L/min, the odor dilution might flow upstream and contaminate the MFC. To avoid these problem, one must first establish the equivalent resistance of the system for both the clean air and the odorized air branches. Then, using shuttle valves, which have two output ports, use the second port to establish an analogous resistance to the one of the system with the valve open. The shuttle valve outputs to port 2 when the solenoid is OFF and to port 1 when the solenoid is ON. Port 1 is connected to the olfactometer, while port 2 is connected to the equivalent resistance tubing and then to vacuum.

### System contamination

A major technical problem in odor delivery systems is contamination. After using a specific odor, the system might retain traces of it that are sufficient to induce unexpected results in the next measurement. In our odor delivery system, using a cleaning cycle after each hour of experiments resulted in no detectable contamination for the photoionization detector or for the *Drosophila* larvae antennal lobe. The cleaning cycle consists in connecting a 5L/min massflow controller in the odorized air branch and running it in purge mode for 30 minutes, although shorter times might be sufficient.

### Speed bottlenecks and control tradeoffs

The system has many potential sources of delay. One of them is the switching time of the valves. Valve switching time may vary between 7ms to 45ms. For our NRESEARCH valves (Figure 1-Figure Supplement 1), driving them with relays resulted in response times of 35 ms, while driving them with high-frequency H-bridge drivers (Accuthermo FTX100) resulted in response times of 10 to 14 ms. Another source of delay is the amount of air flow input needed by the odor dilution bottle to produce odorized air at the output. In this case, the smaller the space in the bottle the faster the input is transformed in output but also the easier it is to saturate the output with smaller airflow values.

The main speed bottleneck in our system is determined by the time it takes for the input signal to produce changes at the output. Despite of our improvement of valve times and the odor bottle delays, the system accumulates around 0.2s of delay between input and output.This results in a speed constraint for precise odor waveform control and it is particularly noticeable for non-continuous odor waveforms like step functions (Ext. Fig. 1A).

### Faster signals in open loop

To produce fast signals, such as white noise, our odor delivery system has to operate in open loop, without the 2-DOF-PID controller, and only using the MFCs PID controller. Using two parallel MFCs with a double shuttle valve further increases the speed of the system (Ext. Fig. 1B left). In this configuration, the time it takes for the MFC to reach a new airflow value is not the bottleneck because the shuttle valves switch between MFCs such that only stable odorized airflows are delivered (Ext. Fig. 1B right). The result is a white process at 40Hz, with flat power spectral density and small auto-correlation times (Ext. Fig. 1C).

---

**Figure 1–Figure supplement 1.**
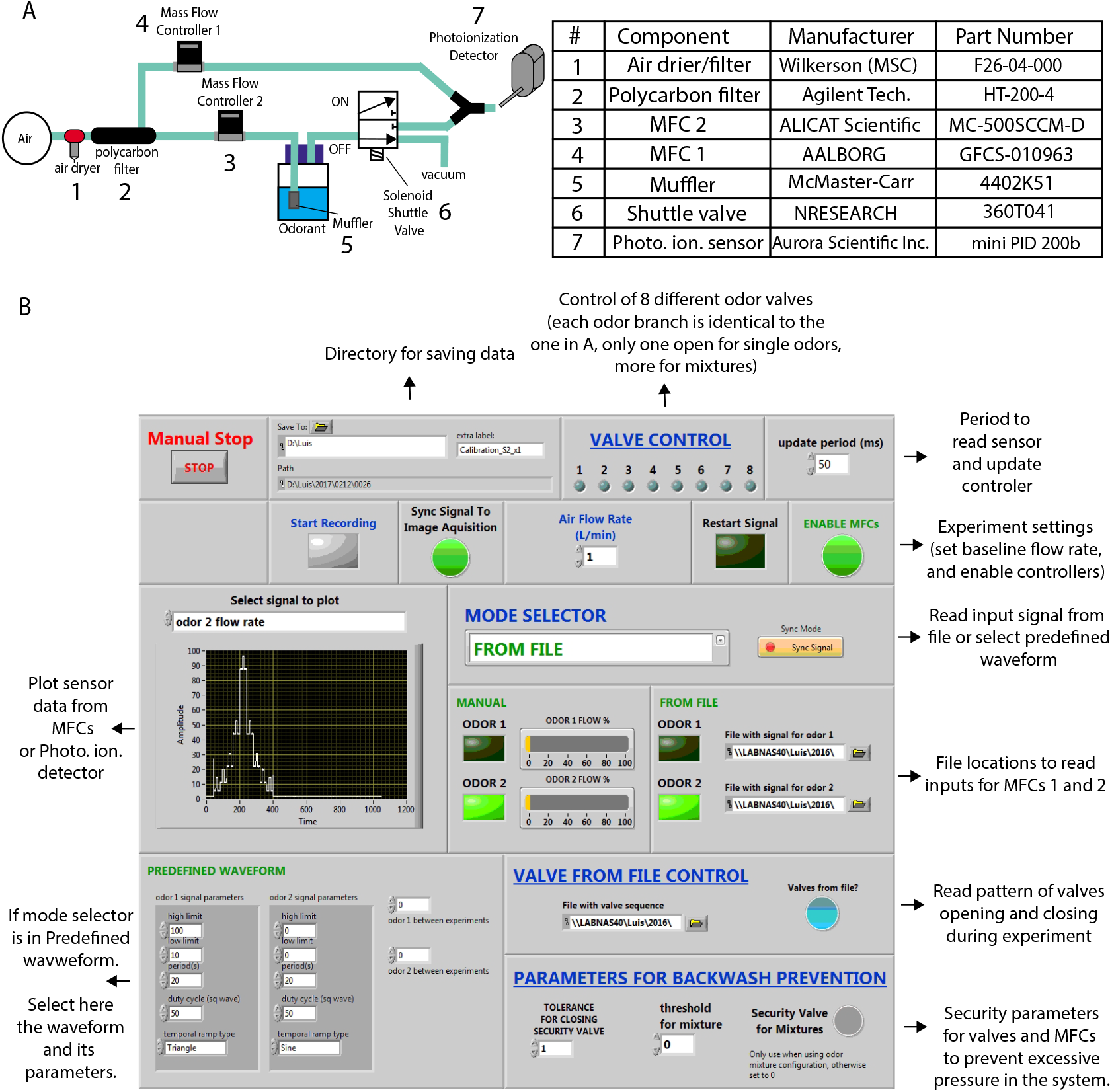
Figure1-Figure Supplement 1. Olfactometer components (**A**) and graphical user interface (GUI) (**B**).

**Figure 4–Figure supplement 1.**
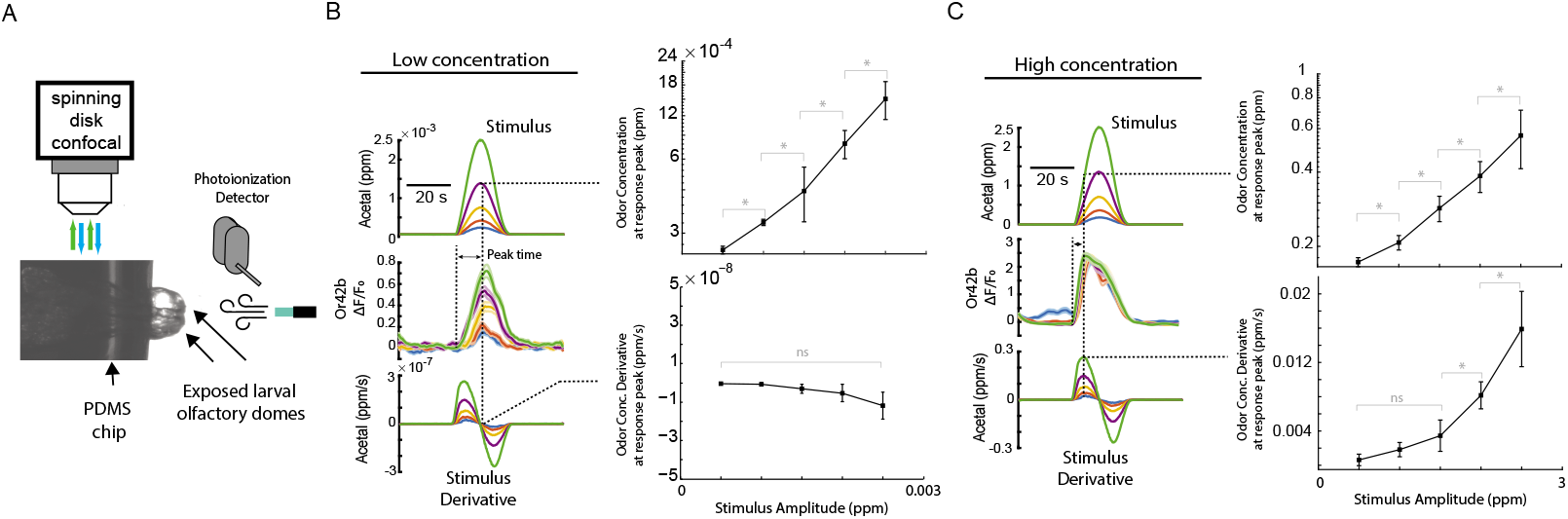
Odor concentration and rate of change at the response peak. **(A)** Experimental preparation for *Drosophila* lavae. Larvae are embedded in a PDMS chip with the olfactory domes exposed and odorized arflow is delivered to the exposed head of the animal. **(B, C)** Odor concentration and derivative at the time point of maximum response amplitude. ‘*’ indicate significant difference with p>0.05 in Wilcoxon matched pairs test. ‘n.s.’ indicates not significantly different. Error bars are the s.e.m.

**Figure 4–Figure supplement 2.**
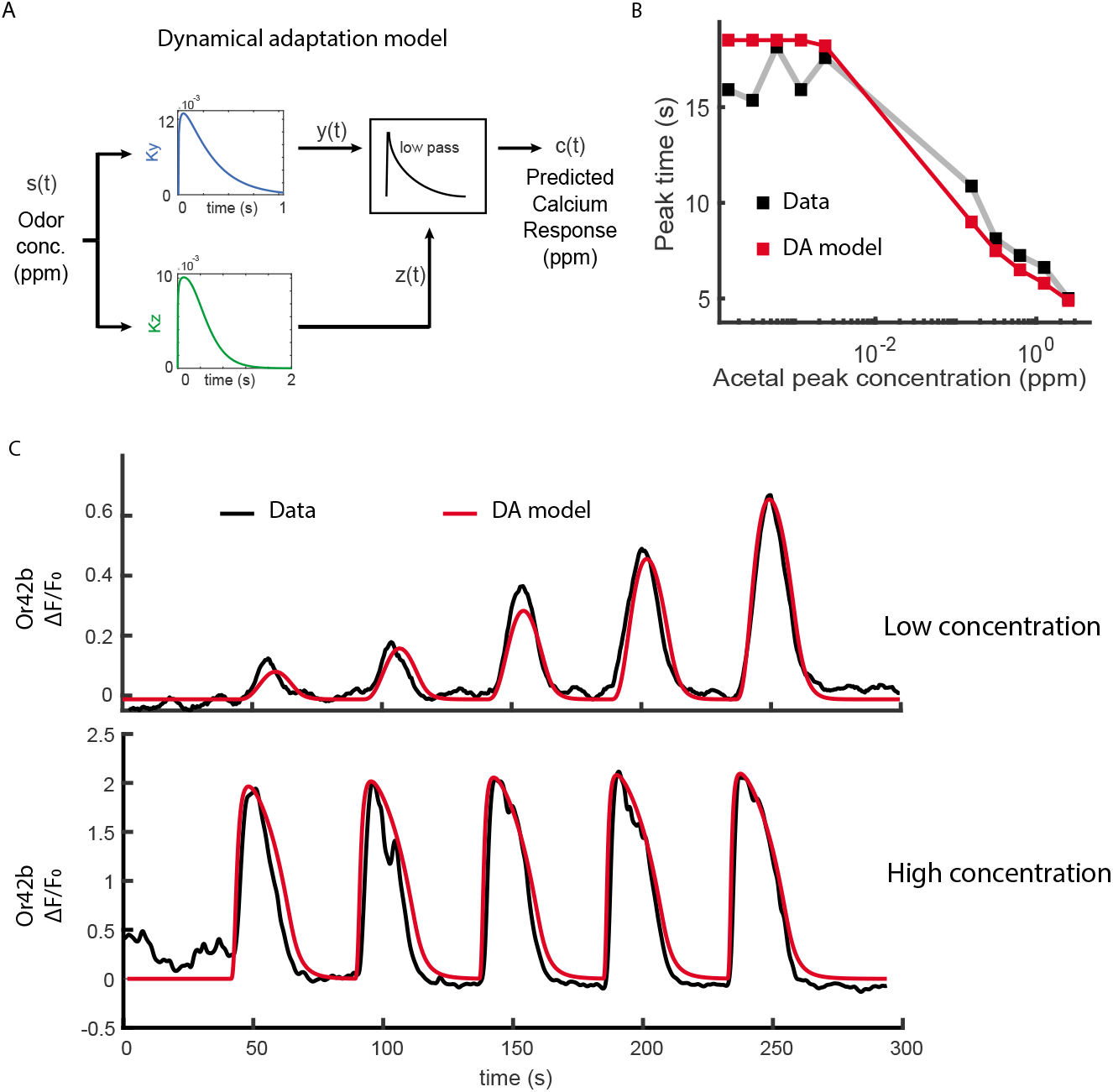
Dynamic Adaptation Model. **(A)** Dynamic Adaptation Model diagram. **(B)** Predicted response peak times match experimental results. **(C)** Predicted responses in red vs. experimental measurements in black.

**Figure 5–Figure supplement 1.**
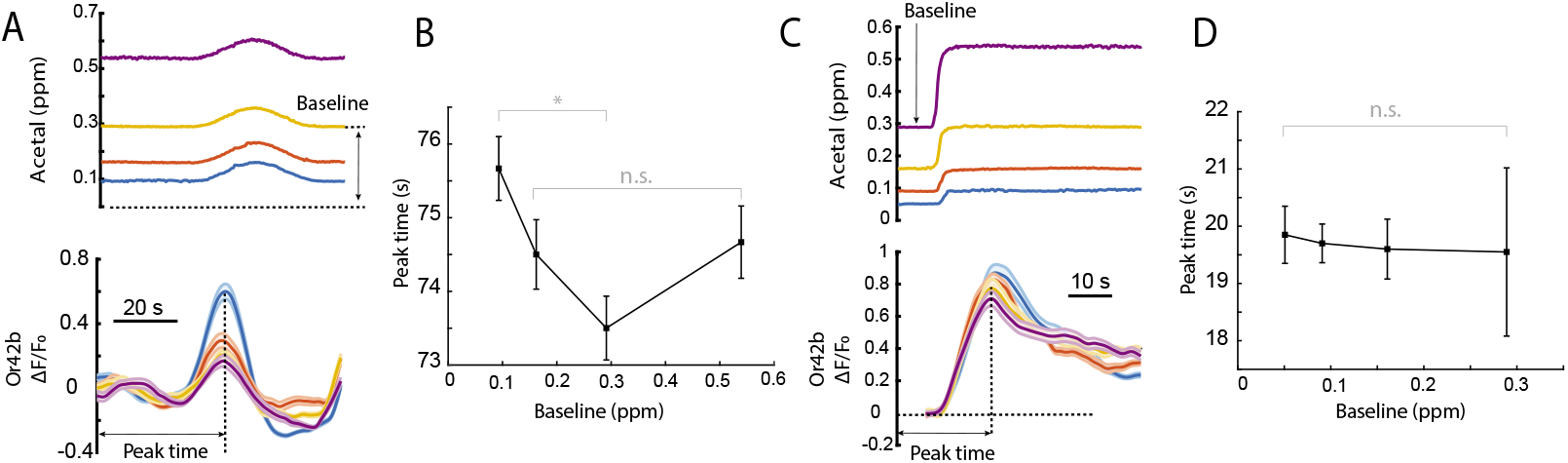
Response peak time vs stimulus baseline. **(A)** Color-coded responses. **(B)** Response peak time vs. stimulus baseline. At lower baselines, responses are faster if amplitude is constant. **(C)** Color-coded responses. **(D)** Response peak time vs. stimulus baseline. If contrast is kept constant, response peak time is independent of stimulus baseline.

**Figure 6–Figure supplement 1.**
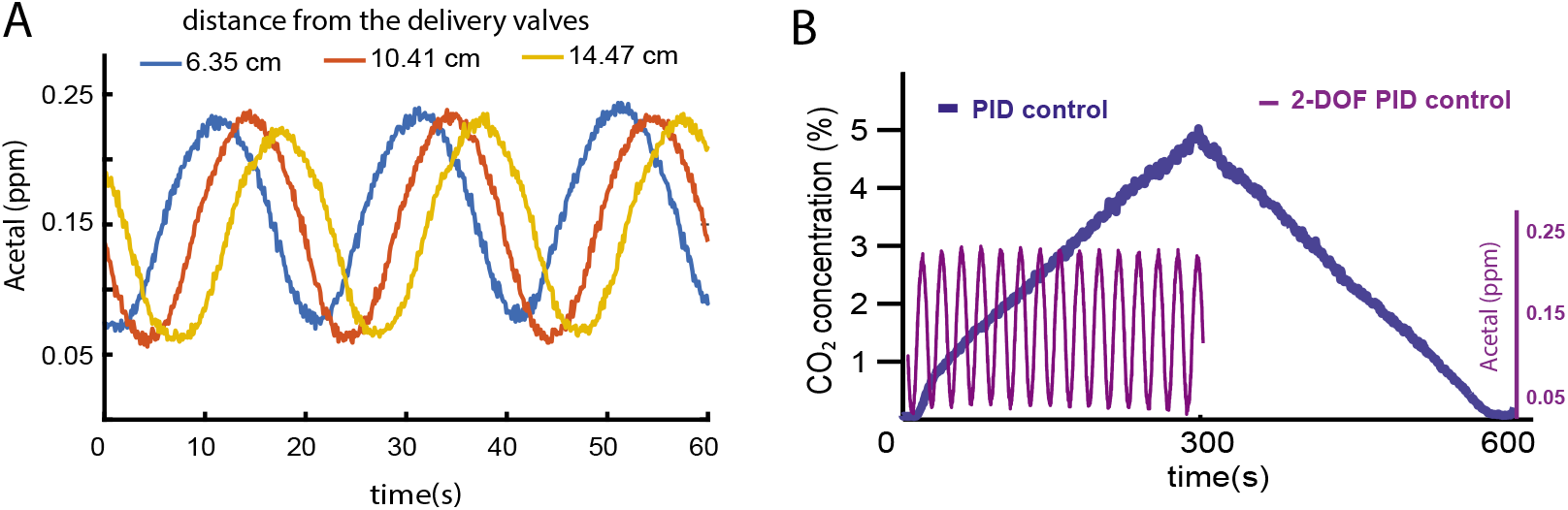
Odor waveforms in a behavioral chamber. **(A)** At different distances of the delivery valves, identical odor waveforms have increasing latencies. **B** 2-DOF-PID controllers enable the delivery of faster odor waveforms than 1-DOF-PID controllers.

**Figure 7–Figure supplement 1.**
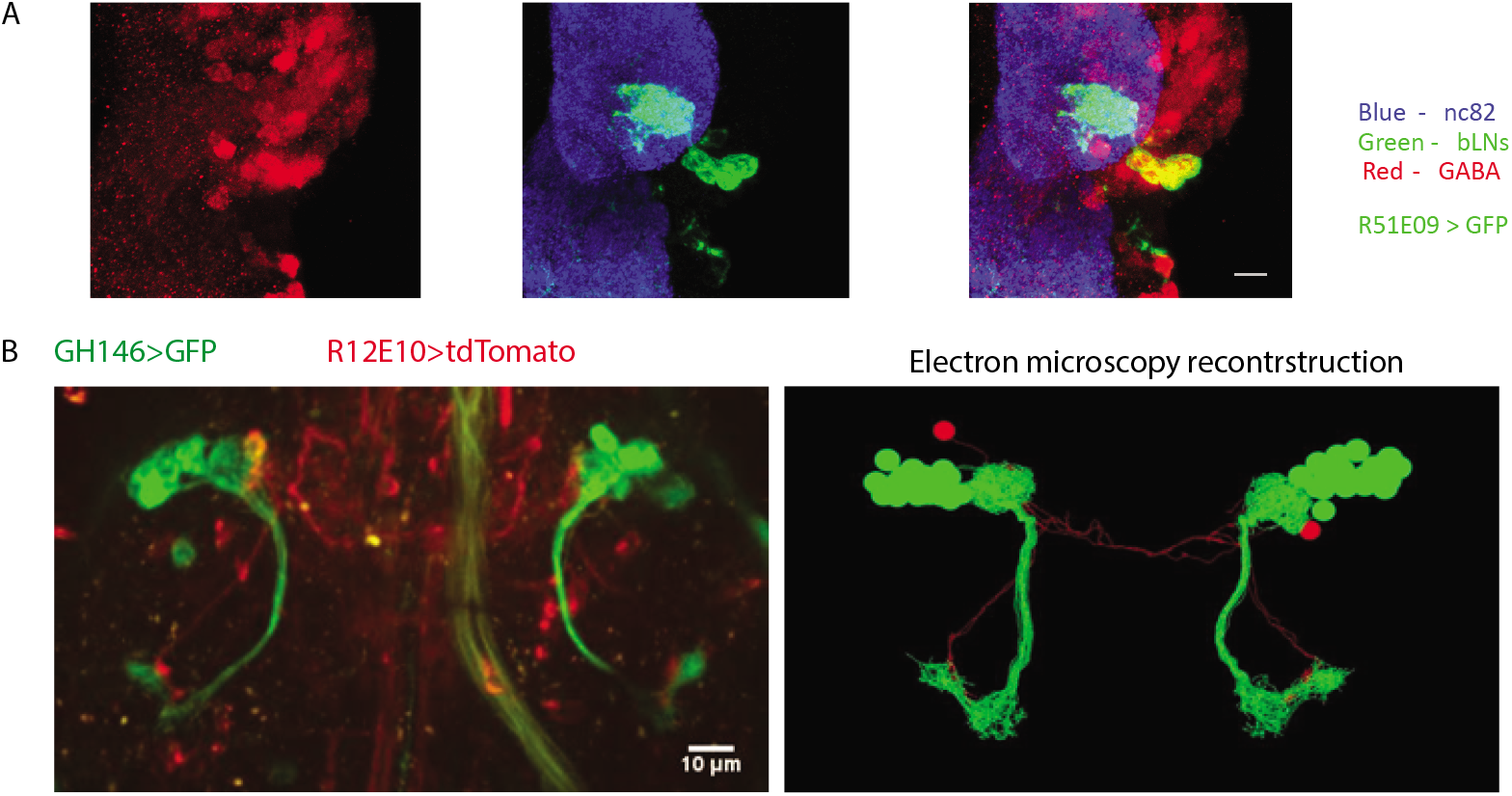
Identification of uniglomerular circuit components. **(A)** Antibody staining with Alexa 488 anti-GFP, Alexa 555 anti-GABA in R51E09-Gal4;UAS-GFP first instar larvae. nc82 was used as a reference. **(B)** Light microscopy vs electron microscopy reconstruction of the uPNs. (GH146-Gal4/LexAop-RFP;R12E10-LexA/UAS-GFP)

